# Combined single-cell profiling of expression and DNA methylation reveals splicing regulation and heterogeneity

**DOI:** 10.1101/328138

**Authors:** Stephanie M. Linker, Lara Urban, Stephen Clark, Mariya Chhatriwala, Shradha Amatya, Davis J. McCarthy, Ingo Ebersberger, Ludovic Vallier, Wolf Reik, Oliver Stegle, Marc Jan Bonder

**Author notes:** shared first authors (Stephanie M. Linker, Lara Urban).

## Abstract

**Background:** Alternative splicing is a key regulatory mechanism in eukaryotic cells and increases the effective number of functionally distinct gene products. Using bulk RNA sequencing, splicing variation has been studied across human tissues and in genetically diverse populations. This has identified disease-relevant splicing events, as well as associations between splicing and genomic variations, including sequence composition and conservation. However, variability in splicing between single cells from the same tissue or cell type and its determinants remain poorly understood.

**Results:** We applied parallel DNA methylation and transcriptome sequencing to differentiating human induced pluripotent stem cells to characterize splicing variation (exon skipping) and its determinants. Our results shows that variation in single-cell splicing can be accurately predicted based on local sequence composition and genomic features. We observe moderate but consistent contributions from local DNA methylation profiles to splicing variation across cells. A combined model that is built based on sequence as well as DNA methylation information accurately predicts different splicing modes of individual cassette exons (AUC=0.85). These categories include the conventional inclusion and exclusion patterns, but also more subtle modes of cell-to-cell variation in splicing. Finally, we identified and characterized associations between DNA methylation and splicing changes during cell differentiation.

**Conclusions:** Our study yields new insights into alternative splicing at the single-cell level and reveals a previously underappreciated link between DNA methylation variation and splicing.

## Background

RNA splicing enables efficient gene encoding and contributes to gene expression variation by alternative exon usage [1]. Alternative splicing is pervasive and affects more than 95% of human genes [2]. Splicing is known to be regulated in a tissue-specific manner [3, 4] and alternative splicing events have been implicated in human diseases [5]. Bulk RNA sequencing (RNA-seq) of human tissues and cell lines has been applied to identify and quantify different splicing events [6], where in particular exon skipping at cassette exons, the most prevalent form of alternative splicing [1], has received considerable attention.

Different factors have been linked to splicing of cassette exons, including sequence conservation [7] and local genomic features such as the sequence composition and the length of the exon and flanking introns [5, 8]. Although there is some evidence for a role of DNA methylation in splicing regulation, this relationship is not fully understood and alternative models have been proposed [9–11]. The transcriptional repressor CTCF has been shown to slow down RNA polymerase II (Pol II), resulting in increased exon inclusion rates. By inhibiting CTCF binding, DNA methylation can cause reduced exon inclusion rate [9]. Alternatively, increased DNA methylation of the MeCP2 pathway has been associated to increased exon inclusion rates. MeCP2 recruits histone deacetylases in methylated contexts that wrap the DNA more tightly around the histones. This interplay between MeCP2 and DNA methylation slows down Pol II, thus leading to an increased exon inclusion rate [10]. Finally, HP1, which serves as an adapter between DNA methylation and transcription factors, increases the exon inclusion rate if it is bound upstream of the alternative exon. Binding of HP1 to the alternative exon leads to increased exon skipping [11]. These alternative mechanisms point to a complex regulation of splicing via an interplay between DNA sequence and DNA methylation, both in proximal as well as distal contexts of the alternative exon.

Technological advances in single-cell RNA-seq have enabled studies that started investigating splicing variation at single-cell resolution [8, 12, 13]. We here leverage very recent protocols for parallel sequencing of RNA and bisulfite treated DNA from the same cell (single-cell methylation and transcriptome sequencing; scM&T-seq [14]) to extend the single-cell splicing analysis by accounting for cell specific DNA methylome profiles. We apply our approach to investigate associations between single-cell splicing variation and DNA methylation at two states of human induced pluripotent stem (iPS) cell differentiation.

## Results

### Single-cell splicing variation during endoderm differentiation

We applied parallel single-cell methylation and transcriptome sequencing (scM&T-seq) to differentiating induced pluripotent stem (iPS) cells from one cell line ("joxm_1") of the Human Induced Pluripotent Stem Cell Initiative (HipSci) [15, 16]. We profiled 93 cells from two different cell types, respectively, namely cells in the iPS state (“iPS”) as well as cells following three days of differentiation towards definitive endoderm (“endoderm”). After quality control this resulted in 84 and 57 cells, respectively (Methods), which were used for analysis. In each cell we quantified cassette exon inclusion rates (Methods, **Table S1-S2**). We quantified splicing rates for between 1,386 and 4,917 cassette exons in each cell (minimum coverage of five reads), estimating splicing rates (PSI) as the fraction of reads that include the alternative exon versus the total number of reads at the cassette exon (Methods). Differences in sequencing depth and cell type explained most of the differences in the number of quantified splicing events between cells (**Figure S1, Table S1-S2**). DNA methylation profiles were imputed using DeepCpG [17], yielding on average 23.1M CpG sites in iPS and 21.6M CpG sites in endoderm cells. We considered 6,265 (iPS) and 3,873 (endoderm) cassette exons that were detected in at least ten cells for further analysis.

Initially, we explored whether individual cells express only a single splice isoform ("cell model", Methods), or whether multiple isoforms are present in a given cell ("gene model"; Methods, **Fig. 1a**), a question that has previously been investigated in bulk and single-cell data [18, 19]. Specifically, we compared the observed distribution of splicing rates PSI in our data to the expectation of a binomial distribution according to the cell model [18], and to the expected distribution according to the gene model (Methods, **Fig. 1a**). Globally, our data rule out the cell model, however we also observed deviations from the gene model, in particular for exons with intermediate levels of splicing (0.2 < PSI < 0.8, **Fig. 1b**).

**Figure 1.**
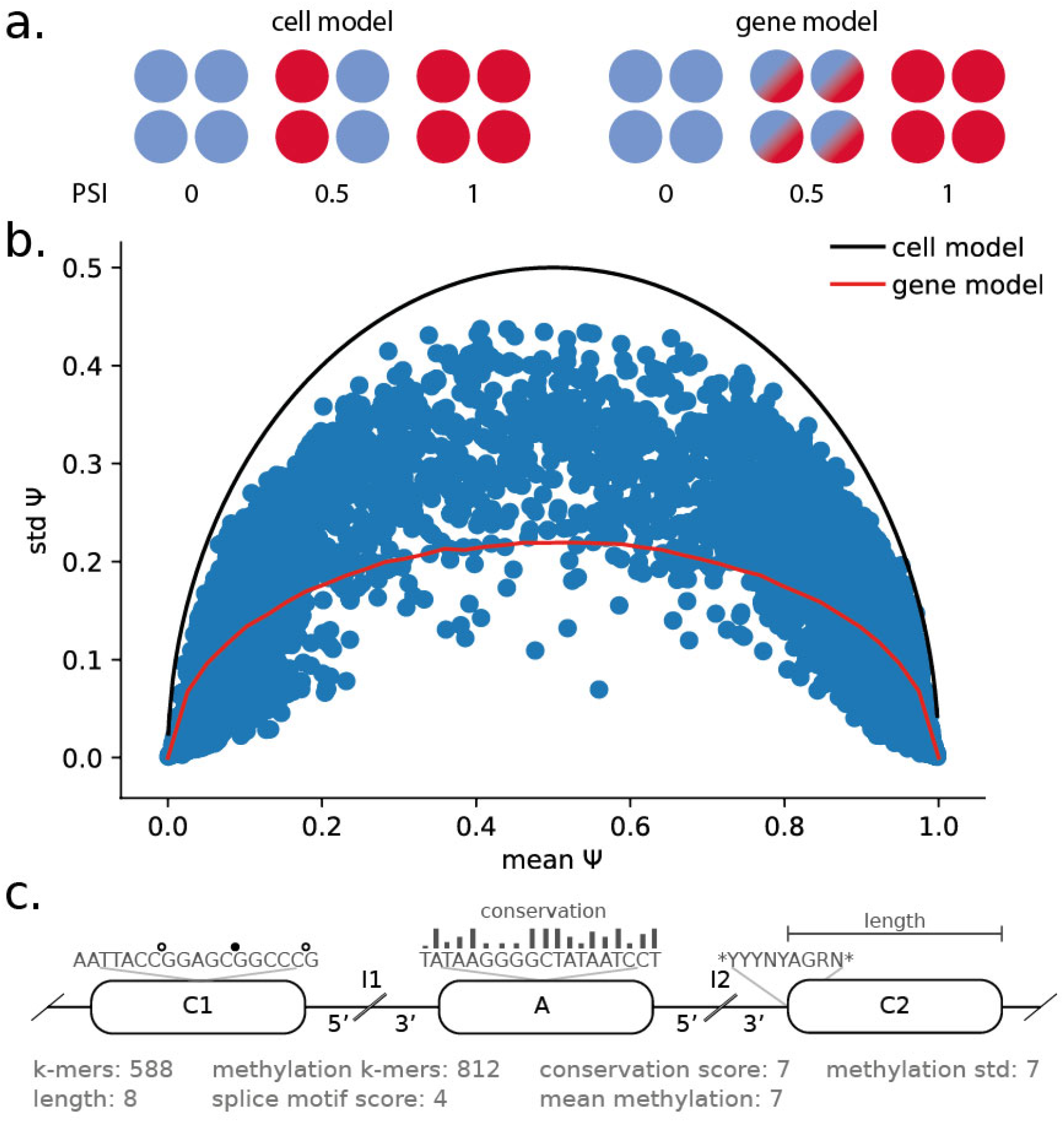
Single-cell splicing and considered features for modeling splicing rates. **a.** Illustration of two canonical splicing models. The "cell model" assumes that splicing variation is due to differential splicing between cells, where each cell expresses only one of two splice isoforms. The "gene model" corresponds to the assumption that both splice isoforms can be expressed in the same cells. **b.** Mean-variance relationships of splicing rates in iPS cells. Shown is the standard deviation of splicing rates across cells for the same cassette exon (standard deviation of PSI) as a function of the average inclusion rate of the cassette exons across cells, considering 84 iPS cells. Solid lines correspond to the expected relationship when either assuming a "cell model" (black line) or when assuming the "gene model" (red line). **c.** Illustration of the considered features and genomics contexts for predicting splicing variation. “A” denotes the alternative exon, “I1” and “I2” correspond to the upstream and downstream flanking introns and “C1” and “C2” to the upstream and downstream flanking exons, respectively. The 5’ and 3’ ends (300 bp) of the flanking introns are considered separately.

### Methylation heterogeneity across cells is associated with splicing variability

Next, to identify locus-specific correlations between DNA methylation heterogeneity and variation in splicing across cells, we tested for associations between differences in imputed DNA methylation levels across cells and splicing rates (Spearman correlation; Methods). For each cassette exon, we tested for associations between splicing rate (PSI) and variation in DNA methylation in each of seven sequence contexts: the upstream, alternative and downstream exons, and the 5’ and 3’ end of the two introns (Methods, **Fig. 1c**). Genome-wide, this identified 424 cassette exons with a methylation-splicing associations in iPS cells (out of 5,564 tested cassette exons, Q < 0.05, **Figure S2a, Table S3**) and 245 associations in endoderm cells (out of 2,811 tested, Q < 0.05, **Figure S2a, Table S3**). The majority of these associations was observed in the upstream alternative exon (~75%), with approximately equal numbers of positive (increased DNA methylation is linked to increased alternative exon inclusion) and negative effects (increased DNA methylation is linked to decreased alternative exon inclusion) (58% of effects as positive in iPS, 55% as positive in endoderm cells). Most associations could be detected in more than one context for a given exon with consistent effect directions (**Figures S2b, c**). Similarly, we observed largely concordant associations across the two cell types in our data. Among exons that are expressed in both iPS and endoderm (n=3,743) we observe that 77% of the associations identified in iPS can be replicated in endoderm cells (P<0.05, with a consistent effect direction), and 89% of the associations identified in endoderm can be replicated in iPS cells (P<0.05, with a consistent effect direction). Genes with negative associations between DNA methylation in the three upstream regions and PSI were enriched for HOXA2 transcription factor binding sites (iPS: 78/118 query genes linked to HOXA2, adjusted P=6.02×10^-4^; endoderm: 60/90 query genes linked to HOXA2, adjusted P=9.03×10^-3^; enrichment based on g:Profiler [20]).

### Prediction of splicing at single-cell level

To gain insights into the global determinants of splicing, we trained alternative regression models to predict splicing rates in single cells using local genomic and epigenetic features (**Fig. 1c**). Briefly, for each cell type we combined splicing rates across all cassette exons and cells and trained a global regression model (assessed using 10-fold cross validation; Methods). Initially, we considered models based on a set of 607 “genomic” features derived from local sequence composition (based on k-mers), sequence conservation and the length of the seven sequence contexts of each cassette exon ("genomic" features, Methods, **Table S4**). Notably, this model yielded a performance that was similar to previous approaches to predict splicing rates using bulk [5] and single-cell [8] RNA-seq (r*2*=0.704, r*2*=0.668; assessed using 10-fold cross validation (CV); **Fig. 2a**, **Figure S3**). To facilitate the comparison with previous results using bulk RNA-seq, we also considered predicting aggregate splicing rates across cells (“pseudo bulk PSI”, bPSI), which resulted in similar prediction accuracies (r2=0.745 and r2=0.733 for iPS and endoderm cells, **Figure S4**).

**Figure 2.**
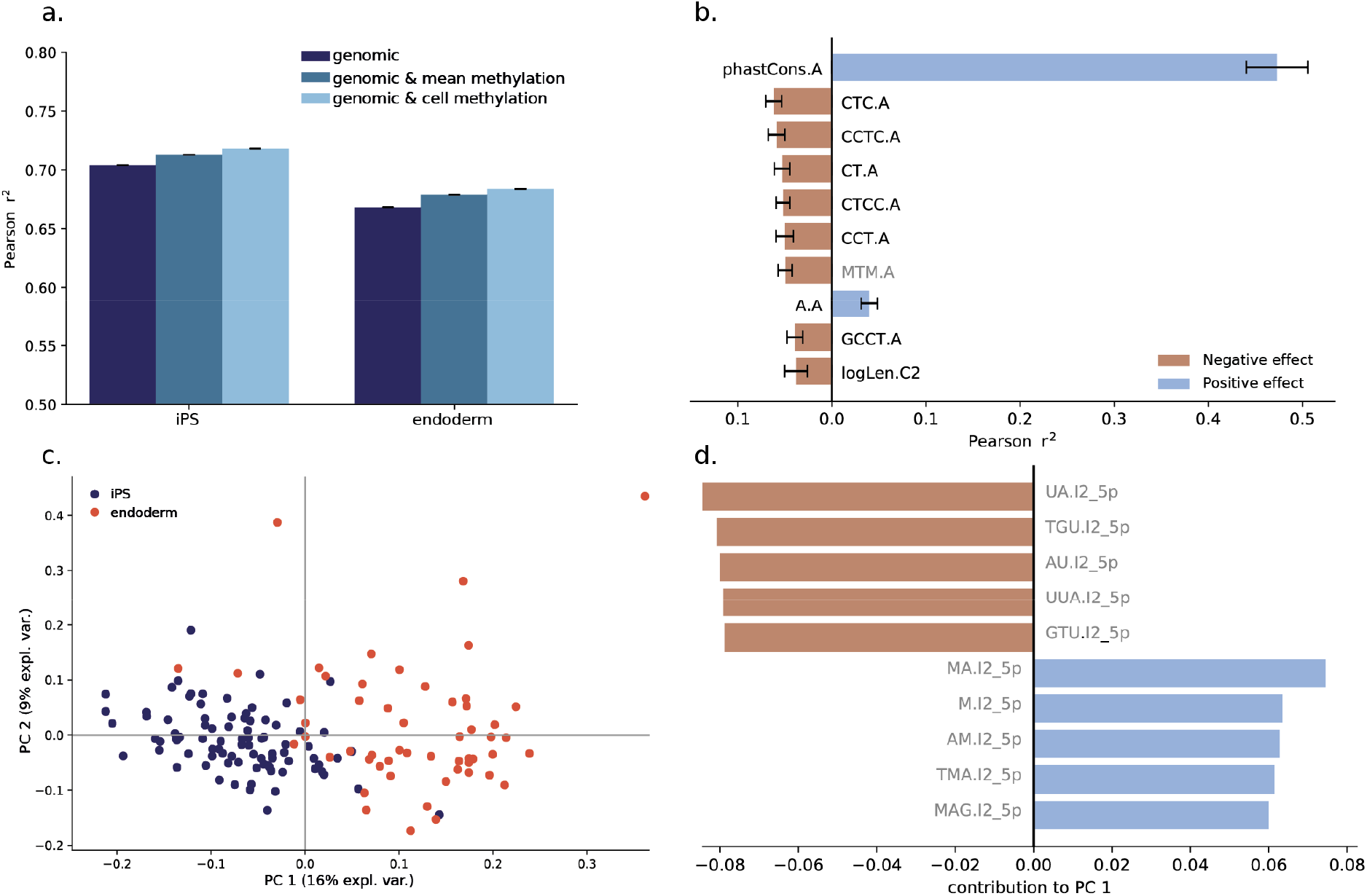
Regression-based prediction of single-cell splicing variation. **a.**Prediction accuracy of alternative regression models for predicting splicing rates in single cells. Shown are out of sample r2 (10-fold CV) in iPS cells and endoderm cells. The genomic model (genomic, dark blue) predicts splicing rates based on sequence k-mers, conservation scores and the length of local contexts (size of the cassette exon, length of flanking introns). Other models account for average methylation rates across cells (genomic & mean methylation, blue), or cell-specific methylation rates (genomic & cell methylation, light blue) as additional features. Error bars denote plus or minus one standard deviation across four repeat experiments. **b.** Relevance of individual features for predicting splicing rates. Shown are correlation coefficients between individual features and PSI values. Features are ranked according to absolute correlation coefficient. The most correlated features are features of the alternative exon (marked “.A”), and include a methylated k-mer (grey). Error bars denote plus or minus one standard deviation of the feature relevance across cells. Positive correlations denote features that are associated with increased inclusion rates of the alternative exon. **c.** Principal component analysis on the feature relevance profiles as in **b** across all cells. The first principal component (PC) primarily captures differences between cell types. **d.** Ten features with the largest contribution to the first PC (five positive and five negative features), which include k-mers with methylation information of the downstream intron I2. Methylation features are shown in grey.

Next, we considered extended models using an additional set of up to 826 DNA methylation features, including an extended k-mer alphabet that takes the methylation status into account, as well as the DNA methylation rate and variance (across CpG sites) in each of seven sequence contexts of a cassette exon (Methods). Methylation features describe methylation patterns of either individual cells ("genomic & cell methylation" features) or averaged across cells ("genomic & mean methylation" features; **Table S4**, **Fig. 1c**). Inclusion of DNA methylation features into the model yielded increased prediction accuracy, where the largest gains were observed when including cell-specific DNA methylation information (“genomic & cell methylation” versus “genomic & mean methylation”). Notably, the inclusion of DNA methylation features when modeling the average splicing rate (“pseudo bulk” models) did not improve the models (**Figure S4**). This observation in combination with our previous results suggests that DNA methylation is most predictive of cell-to-cell variation in splicing in a locus-specific manner, whereas genomic features by design can only explain variation across different loci. These findings were consistent across both cell types, and we observed analogous benefits of accounting for DNA methylation when applying the same models to a previous scMT-seq datasets from mouse ES cells [14] (Methods, **Figures S3, S4**).

Next, to assess the relevance of the the considered features, we considered regression models based on individual features trained in individual cells. Consistent with previous bulk studies [5, 7], this identified features derived from the alternative exon and its neighboring contexts, namely the 3’ end of the upstream intron and the 5’ end of the downstream intron, as most informative (**Table S5**). Within these contexts, sequence conservation of the alternative exon was identified as the most relevant feature. Other relevant features included the k-mers CT, CTC and CCT of the alternative exon (**Fig. 2b)**, sequence patterns that show close resemblance to CTCF binding motifs, which has previously been linked to alternative splicing. Notably, unlike previously described CTCF or CTCF-like motifs that are located upstream [9] or downstream [21] of the alternative exon and increase inclusion rate of the alternative exon, these k-mers are located in the alternative exon and decrease the inclusion rate [9, 21].

The relevance of the cell-specific features for splicing prediction as quantified by regression weights were highly consistent across iPS and endoderm cells. This consistency extended to the mouse embryonic stem (ES) cell dataset, where again the features of the alternative exon and sequence conservation scores were the most relevant predictors for splicing (**Table S5, Figure S5**). Despite the overall consistency in feature relevance (r^2^=0.79, average correlation between weights across all iPS and endoderm cells), principal component analysis (PCA) applied to the feature relevance identified subtle coordinated differences of the feature relevance between cell types (**Fig. 2c**). The first two principal components (PC) clearly separate iPS from endoderm cells, differences that are primarily attributed to k-mers of the downstream intron (I2) that contain methylated and unmethylated cytosine bases (**Fig. 2d**, **Table S6**). Consistent with this, a single-cell methylation model trained on endoderm cells yielded only moderate prediction accuracy in iPS cells (r2=0.52), highlighting the cell type specificity of the models that include DNA methylation information. This points towards a combination of differences in sequence composition, potentially transcription factor activity, and DNA methylation as the main determinants of cell-type specific splicing regulation.

Finally, we also considered more complex regression models based on convolutional neural networks to predict single-cell splicing based on DNA sequence and an extended genomics alphabet including base-level DNA methylation information (deposited at kipoi [22], Methods). We observed only a limited increase in performance when including DNA methylation information (**Supplementary Results**, **Figure S6**). These results line up with the locus-specific DNA methylation and the linear regression results, supporting the hypothesis that global splicing information is primarily encoded by DNA sequence and conservation, and DNA methylation is linked to splicing in a locus-specific manner.

### Prediction of splicing modes of individual exons

Next, we set out to study differences between different exons and their splicing patterns. We classified cassette exons into five distinct categories, using a scheme similar to that of Song *et al.* [12]: 1) excluded, 2) included, and three intermediate splicing categories: 3) overdispersed, 4) underdispersed and 5) multimodal (**Fig. 3a, 3b, Table S7**, Methods). We trained and cross-validated multinomial regression models (Methods) to classify individual exons using analogous feature sets as considered for the regression models on single-cell splicing (**Table S4**). A model based on genomic features yielded a macro-average AUC of 0.85 in iPS (**Fig. 3c**) and 0.84 in endoderm (**Figure S7**), where again sequence conservation in different contexts was the most informative feature (**Table S8**). Interestingly, we observed differences in the feature relevance across splicing categories: i) included and excluded exons, where the most relevant features were located in the alternative exon, and ii) the intermediate splicing categories, where features of the flanking exons were most informative. In general, predictions of included and excluded exons were most accurate (AUC=0.96 for both in iPS, AUC=0.94 for included in endoderm, AUC=0.96 for excluded in endoderm cells, **Fig. 3d**, **Figure S7a**). These prediction accuracies exceed previously reported results in bulk data [5]. Even higher accuracies were achieved when training a model to discriminate between included and excluded exons only (AUC=0.99 in iPS), whereas lower prediction accuracies were achieved for discriminating the intermediate splicing categories only, both in iPS and endoderm cells (AUC=0.7-0.9, **Table S8**). The inclusion of the DNA methylation features did not improve the prediction performance (**Fig. 3d**, **Figure S8a**).

**Figure 3.**
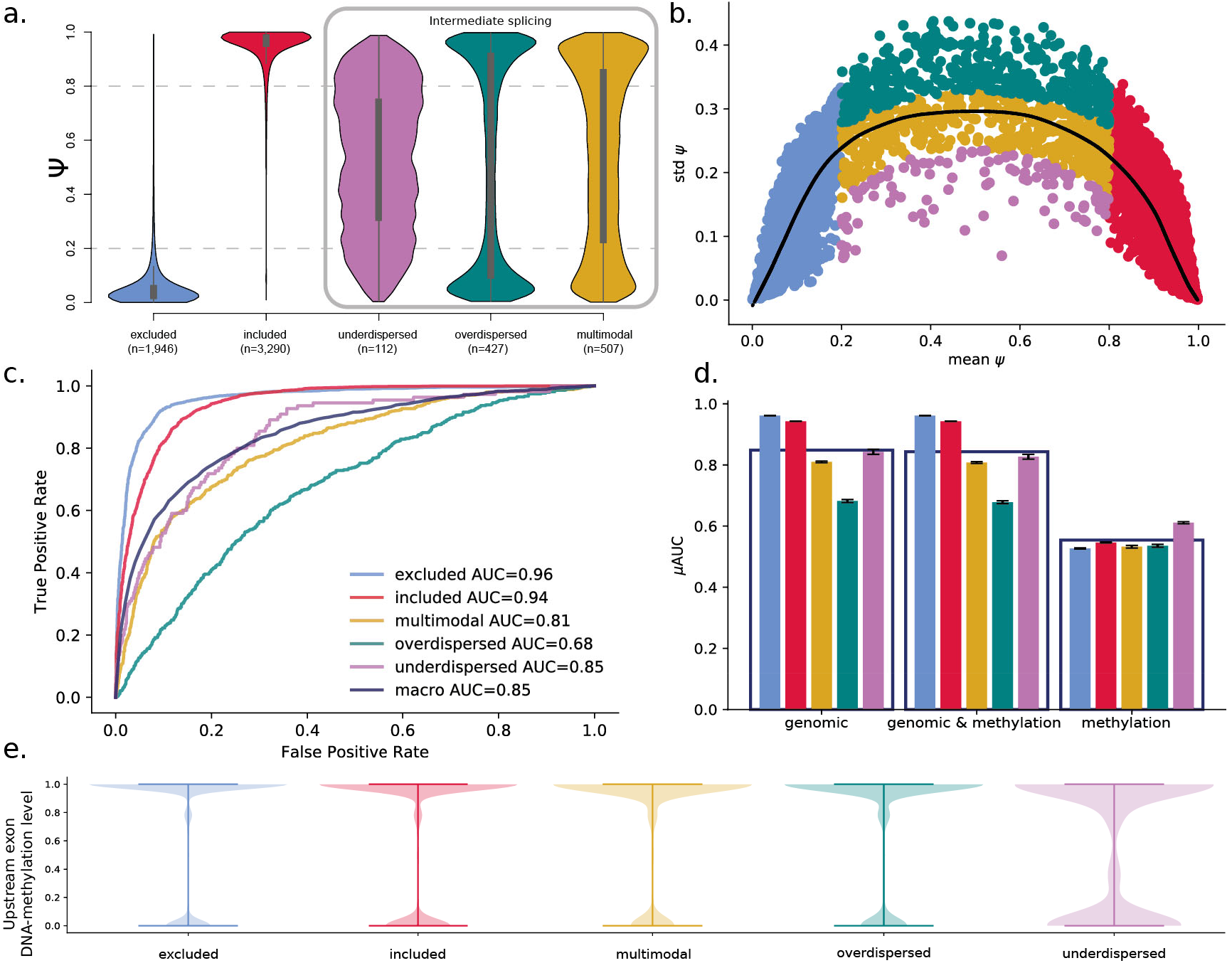
Classification of cassette exons based on single-cell splicing patterns in iPS cells. **a.** Single-cell splicing rate (PSI) distributions of the five splicing categories (inspired by Song *et al.* [12]) in 84 iPS cells. The intermediate splicing categories that can only be defined based on single-cell information are framed by a grey box. **b.** Variation of PSI (standard deviation) across cells as a function of the average inclusion rate of cassette exons across 84 iPS cells, colored according to their respective splicing category as defined in **a**. The solid black line denotes the LOESS fit across all cassette exons. **c.** Prediction performance of logistic regression for predicting splicing categories based on genomic features. Shown is the receiver operating characteristics for each splicing category and the macro average (area under the curve, AUC)**. d.** Prediction performance of alternative regression models for each splicing category, either considering a model trained using genomic features (‘genomic’, left), genomic and all DNA methylation features (‘genomic & methylation’, center) as well as only DNA methylation features (‘methylation’, right). The genomic model includes k-mers, conservation scores and region lengths (see Fig. 1c). The genomic and methylation model additionally includes DNA methylation features. The methylation model includes average DNA methylation features per sequence context. Splicing categories are coded in color as in **a**. Error bars denote plus or minus one standard deviation across four repeat experiments. **e**. Distribution of DNA methylation levels in the upstream exon (C1) per splicing category. Methylation is decreased in underdispersed exons.

Consistent with this, we found that a model based on DNA methylation alone did not yield accurate predictions although methylation contained some information for identifying underdispersed cassette exons (**Fig. 3d**, **Figure S8b**). Given this, we investigated the distribution of DNA methylation patterns across splicing categories, observing distinct distributions of DNA methylation in the upstream exon of underdispersed cassette exons (**Fig. 3e**). This effect was consistent, although less pronounced, in other sequence contexts (decreasing from the upstream to the downstream exon, **Figures S9a-b**).

We assessed robustness of these results by cross-replicating between iPS and endoderm cells, and by replicating the results in mouse ES cells. First, we trained the genomic model on endoderm cassette exons and assessed this model’s predictions on iPS-specific cassette exons. The performance of this model was similar to within cell type prediction performance (macro AUC=0.82, **Figure S10a**). However, inclusion of the DNA methylation features into the model resulted in a decline in the cross-prediction performance (macro AUC=0.54, **Figure S10b**). As in the linear model cross-replication analysis, this finding emphasizes the importance of cell type-specific DNA methylation for accurately predicting splicing. Next, the performance for splicing category prediction in mouse ES cells was very similar to the performance in the endoderm and iPS cells (macro AUC=0.82, in the genomic and the genomic and methylation model). We also observed the same distinct distributions of DNA methylation in the upstream exon of underdispersed cassette exons (Figure S9c). However, the relationship between the DNA methylation levels and underdispersed cassette exons category could not be replicated in the mouse cells (**Figure S7b**).

### Splicing category switches across cell differentiation

Finally, we assessed changes in the splicing category switches between cell types. Similar to previous observations in the context of neuronal iPS differentiation [12], we observed that a majority (88%) of the cassette exons retained their category during differentiation (**Fig. 4a**). We also observed no cassette exon that switched from included to excluded or *vice versa.* Instead, most (55%) of the switching events were observed within the three intermediate splicing categories. The most prevalent switch events were changes to the multimodal category; 51% of the underdispersed and nearly 45% of the overdispersed cassette exons in iPS cells switched to multimodal at the endoderm state.

**Figure 4.**
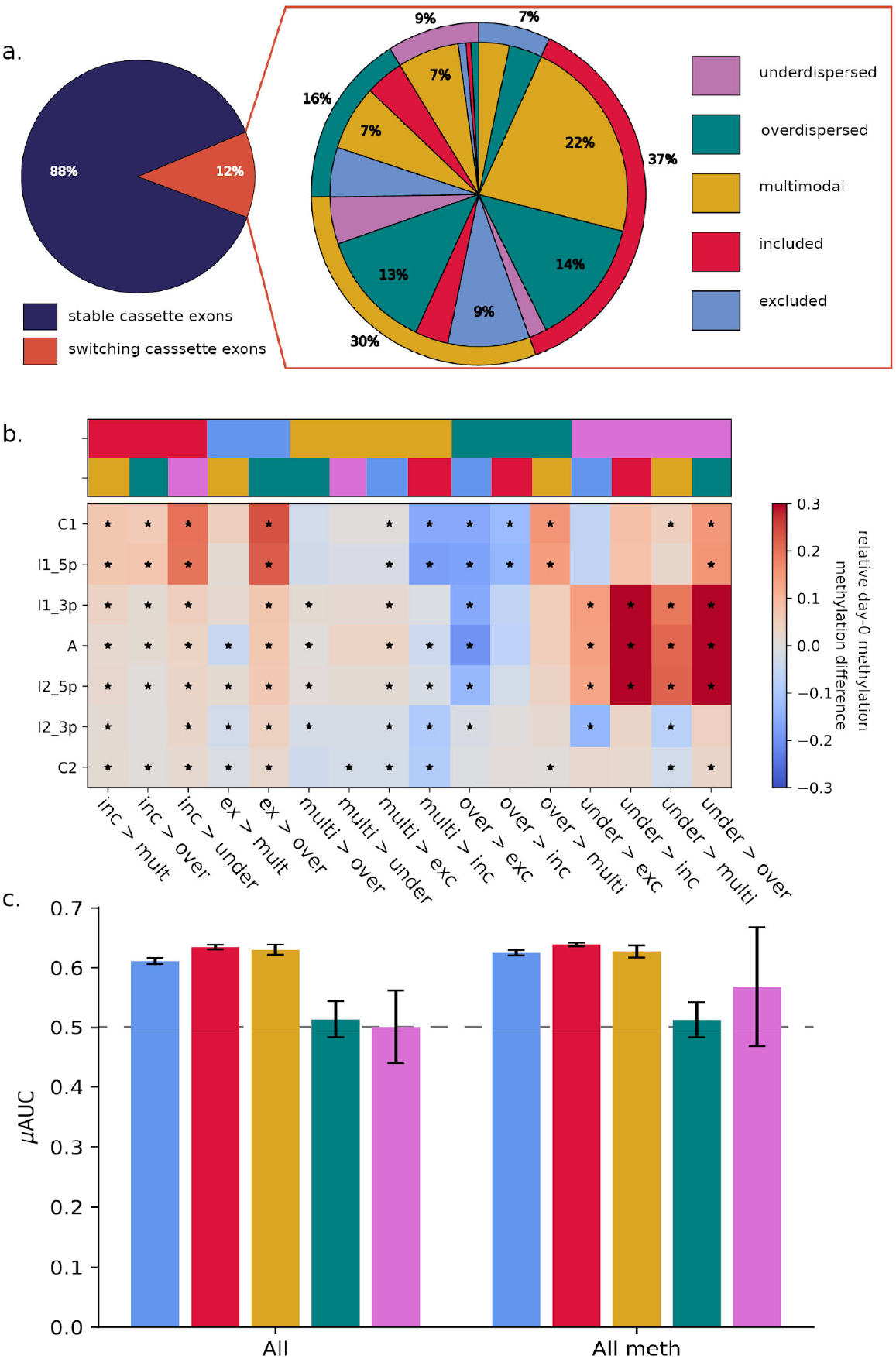
Comparison of splicing category distributions between iPS and endoderm cells. **a.** Pie chart showing the number of category switches between iPS and endoderm cells (left panel). The zoom-in (right panel) shows details of different category switches. The outer pie chart shows the splicing category of each cassette exon at the iPS state and the internal pie chart shows the respective category at endoderm state. Non-annotated slices in the pie chart reflect ~1% of the data. **b.** DNA methylation changes associated with the observed category switches. The top panel shows the iPS and endoderm splicing categories colored according to **a**. The bottom panel shows DNA methylation levels within the seven sequence contexts of a cassette exon as compared to the DNA methylation levels of the cassette exons that do not switch in their splicing category. Significant changes (Q < 0.05) are marked with a star. DNA methylation of the alternative exon and its vicinity is increased in cassette exons that switch from the underdispersed category. Cassette exons that switch from either included or excluded to any other splicing category show increased DNA methylation of the upstream exon (C1). **c.** Performance of logistic ridge regression models that predict absence/presence of switching splicing categories between iPS and endoderm states. DNA methylation information improves prediction of the under- and overdispersed cassette exons. The categories are colored according to **a**. Error bars denote plus or minus one standard deviation across four repeat experiments.

After observing the category switches between the cell types, we set out to build a final set of logistic ridge regression models based on genomic and methylation features to predict category switching ability of cassette exons during differentiation (**Fig. 4c** for prediction performance, **Table S9**). This model had limited power to predict category switches, and DNA methylation did not significantly improve the prediction of any category although moderately higher predictions can be seen for the switching behavior of over- and underdispersed cassette exons.

Lastly, we assessed if DNA methylation changed within the cassette exons switching between the cell types. The DNA methylation levels of cassette exons that switched category only changed minimally (**Figure S11**). However, we observed that DNA methylation of the alternative exon of switching cassette exons differed from non-switching cassette exons at the iPS state (**Fig. 4b**). DNA methylation of both, switching included and switching excluded cassette exons, was increased around C1 in comparison to their relevant non-switching counterparts. In the case of switching overdispersed cassette exons, we observed higher DNA methylation levels within and in the vicinity of the alternative exon.

## Discussion

Here, we present the first analysis of alternative splicing in single cells that considers both genomic and epigenetic factors. Our study focuses on variation of splicing in cassette exons at two different states of cell differentiation. We show that splicing events do not strictly follow the previously suggested cell or gene models of splicing patterns, but instead we find a substantial proportion of exons that are better described by an intermediate model (**Fig. 1b**).

We show that single-cell splicing of cassette exons is influenced by genomic features as previously assessed in bulk data, but also by DNA methylation differences. We observe that DNA methylation is related to splicing phenotypes, with the strongest link to single-cell splicing ratios. When assessing splicing variation in bulk populations (“pseudo bulk”), most of the information encoded in DNA methylation is lost. A reason for this might be the strong correlation between genomic and methylation features, in particular between DNA methylation and cytosine-related features. Additionally, our results suggest that the relationship between splicing and DNA methylation is locus-specific (**Figure S2**). This might explain the lack of prediction performance increase when DNA methylation is included in across-cell models.

Besides sequence conservation, a feature that has previously been described in bulk studies [7], the most relevant features to predict splicing were the k-mers CTC, CT and CCT within the alternative exon (**Fig. 2b**). These k-mers point towards involvement of CTCF. Previous work has shown that CTCF motifs within introns are linked to splicing by slowing down RNA Polymerase II, thereby leading to a higher chance of exon inclusion [9]. Interestingly, there is a known link between DNA methylation and CTCF motifs [9]. Methylation of CTCF binding sites can block CTCF and thereby result in decreased inclusion rates of an exon. As the methylated k-mer equivalents were less predictive of splicing, we suggest a more complex involvement of DNA methylation in alternative splicing, potentially by locus-specific effects, which our current models are not able to capture.

In addition to modelling splicing ratios, we also modelled splicing categories to gain insights into variability of splicing across single cells (**Fig. 3**). The categories considered in our model reflect both, the overall splicing rate and splicing variability across cells. Exons with included versus excluded splicing states could be accurately predicted. In contrast, the intermediate splicing categories which are reflective of single-cell variability could only be predicted with a lower accuracy. This might be due to the lower number of cassette exons assigned to these categories (multimodal n=506, overdispersed n=427, underdispersed n=110, versus included n=3,278 and excluded n=1,944 in iPS cells), or reflect increased vulnerability to assay noise or more complex regulatory dependencies. As in the linear regression models, we observed that DNA sequence conservation scores were the most informative features for predicting splicing categories (**Table S5**). Interestingly, for intermediate categories, the genomic information in the vicinity of the alternative exon rather than of the exon itself seemed to be predictive of splicing variability. Whereas DNA methylation did not contribute to improving the splicing prediction, we observe that DNA methylation levels of underdispersed cassette exons were significantly reduced in all genomic contexts, most significantly in the upstream exon. We hypothesize that the lower DNA methylation levels of underdispersed cassette exons gives the sequence motifs more power to control splicing levels, i.e. increased DNA methylation levels lead to more stochasticity in splicing. This hypothesis is supported by the effect direction of methylation features, which are opposite between overdispersed and underdispersed cassette exons. We finally observe that the methylation k-mers are on average less informative of splicing than non-methylation features, potentially further supporting our hypothesis.

By leveraging data from two cell types, we were able to show the consistency of splicing prediction and of the relevant genomic and methylation features, and could simultaneously assess splicing category maintenance during cell differentiation (**Fig. 2c**). The differences between features predictive of splicing between iPS and endoderm cells were primarily observed within the (methylated) k-mers, which is consistent with the known alteration of transcription factor activity and DNA methylation differences between cell types. Next, we were able to confirm the findings from Song *et al.* [12] that only a limited number of cassette exons switch splicing categories between cell types (**Fig. 4a**). Additionally, as previously described in context of neural differentiation [12], switches between included and excluded categories were not observed. Most of the category switches were observed within the three intermediate splicing categories. Hereby, DNA methylation differences seemed to predate the switching ability. Using ridge regression, we were able to predict if a cassette exon would switch its splicing category between the cell types. Again, DNA methylation seemed to be particularly informative of intermediate splicing. It improved the predictability of switching in over- and underdispersed categories.

The novelties of our analyses also represent their main limitations. Single-cell sequencing intrinsically delivers fewer reads to assess gene expression and DNA methylation levels. Especially the genome coverage of the bisulfite-treated DNA sequencing remains low due to the low quantities of starting material. Using computational imputation we were able to mitigate this effect to some extent. However, imputation strategies have limitations and in particular loci that lack methylation information cannot be recovered.

The intrinsic properties of single-cell data also affect the accuracy of estimated splicing ratios per cassette exon. We opted for a lenient threshold on read depth to determine splicing ratio, which delivered more cassette exons to train our models, but also rendered splicing ratios less accurate in comparison to deep-sequenced bulk data. The low read depth increases the chance of missing an isoform or cassette exon, an effect known as a dropout. Dropouts in single-cell RNA-seq data can have a strong impact on the fit of the cell- or gene-model. If one of the isoforms was completely unobserved, this would decrease the fit of the gene model. On the contrary, sequencing multiple cells at once would decrease the fit of the cell model. Given that our results are robust across cassette exons, cell types and species, the overall findings we report are however not likely to be affected.

## Conclusions

In summary, we showed for the first time that alternative splicing and splicing variability across cells can be predicted with genomic and DNA methylation information in single cells. We assessed the impact of DNA methylation and cellular features on cassette exon splicing, and were able to replicate our findings in two human cell types and mouse ES cells. We investigated stability and variance of splicing between the two cell types and, importantly, we showed that DNA methylation primes splicing switches during differentiation.

## Methods

Single-cell transcription and methylation data was generated from a single donor from the Human Induced Pluripotent Stem Cells Initiative (HipSci) [15, 16], using the previously described protocol for single-cell methylation and transcriptome sequencing in the same cells (scM&T-seq) (see [14] for details). Line joxm_1, an induced pluripotent stem cell (iPSC) line derived from fibroblasts cells from HipSci project, was cultured and triggered into differentiation towards endoderm. scM&T-seq data was generated for 93 cells (together with one empty well as negative control and two 15-cell and 50-cell positive controls) at the undifferentiated time point (iPS) and the definitive endoderm time point (endoderm), yielding 186 cells for analysis.

### Cell handling and differentiation

The joxm_1 IPSC line was cultured in Essential 8 (E8) media (LifeTech) according to the manufacturer’s instructions. For dissociation and plating, cells were washed 1× with DPBS and dissociated using StemPro Accutase (Life Technologies, A1110501) at 37°C for 3 - 5 min. Colonies were fully dissociated through gentle pipetting. Cells were washed 1× with MEF medium [23] and pelleted gently by centrifuging at 285×g for 5 min. Cells were re-suspended in E8 media, passed through a 40 μm cell strainer, and plated at a density of 60,000 cells per well of a gelatin/MEF coated 12 well plate in the presence of 10 μM Rock inhibitor – Y27632 [10 mM] (Sigma, Cat # Y0503 - 5 mg). Media was replaced with fresh E8 free of Rock inhibitor every 24 hours post plating. Differentiation into definitive endoderm commenced 72 hours post plating as previously described [23].

### FACS preparation and analysis of cells

During all staining steps, cells were protected from light. Cells were dissociated into single cells using accutase and washed 1× with MEF medium as described above. Approximately 1 × 10^6^ cells were resuspended in 0.5 mL of differentiation state specific medium containing 5 μL of 1 mg/mL Hoechst 33342 (Thermo Scientific). Staining with Hoechst was carried out at 37°C for 30 min. Unbound Hoechst dye was removed by washing cells with 5 mL PBS + 2% BSA + 2 mM EDTA (FACS buffer); BSA and PBS were nuclease-free. For staining of cell surface markers Tra-1-60 (BD560380) and CXCR4 (eBioscience 12-9999-42), cells were resuspended in 100 μL of FACS buffer with enough antibodies to stain 1 × 10^6^ cells according to the manufacturer’s instructions, and were placed on ice for 30 min. Cells were washed with 5 mL of FACS buffer, passed through a 35 μM filter to remove clumps, and re-suspended in 250 μL of FACS buffer for live cell sorting on the BD Influx Cell Sorter (BD Biosciences). Live/dead marker 7AAD (eBioscience 00-6993) was added just prior to analysis according to the manufacturer’s instructions and only living cells were considered when determining differentiation capacities. Living cells stained with Hoechst but not Tra-1-60 or CXCR4 were used as gating controls.

### scM&T-seq

As previously described in Angermeuller *et al.* [14], scM&T-seq library preparation was performed following the published protocols for G&T-seq [24] and scBS-seq [25], with minor modifications as follows. G&T-seq washes were performed with 20 μl volumes, reverse transcription and cDNA amplification were performed using the original Smart-seq2 volumes [26] and Nextera XT libraries were generated from 100-400 pg of cDNA, using 1/5 of the published volumes. RNA-seq libraries were sequenced as 96-plexes on a HiSeq 2000 using v4 chemistry and 125 bp paired end reads. BS-seq libraries were sequenced as 24-plexes using the same machine and settings, which yielded a mean of 7.4M raw reads after trimming.

### Gene expression quantification

For single-cell RNA-seq data, adapters were trimmed from reads using Trim Galore! [27–29], using default settings. Trimmed reads were mapped to the human reference genome build 37 using STAR [30] (version: 020201) in two-pass alignment mode, using the defaults proposed by the ENCODE consortium (STAR manual). Expression quantification was performed separately using Salmon [31] (version: 0.8.2), using the “--seqBias”, “--gcBias” and “VBOpt” options on transcripts derived from ENSEMBL 75. Transcript-level expression values were summarized at gene level (estimated counts) and quality control of scRNA-seq data was performed using scater [32]. Cells with the following features were retained for analysis: (i) at least 50,000 counts from endogenous genes, (ii) at least 5,000 genes with non-zero expression, (iii) less than 90% of counts are assigned to the top 100 expressed genes per cell, (iv) less than 20% of counts are assigned to ERCC spike-in sequences, and (v) a Salmon mapping rate of at least 40%. These filters jointly removed 9 iPS cells and 36 endoderm cells from our analysis.

### Splicing quantification

Of the 186 cells, 84 (iPS) and 57 (endoderm) cells passed QC on gene expression data as described above. Exon splicing rates in individual cells were quantified using the data-dependent module of BRIE [8]. BRIE calls splicing at predefined cassette exons and quantifies splicing using exon reads in single-cell data. By default BRIE combines informative prior learned from sequence features and a likelihood calculated from RNA-seq reads by a mixture modelling framework that is similar to MISO [33]. As our aim is to model the local and global determinants of splicing, we used splicing rate estimates based on the observed data at individual exons only. We detected and quantified splicing for between 1,386 and 4,917 exons per cell (minimum coverage five reads, in total considered 6,265 (iPS) and 3,873 (endoderm) cassette exons that were detected in at least ten cells for further analysis.

The following settings were used to quantify splicing with BRIE: exons have to be located on autosomes and input chromosomes and should not be overlapped by any other alternatively spliced exon. The surrounding introns have to be longer than 100 bp, the length of the alternative exon regions has to be between 50 and 450 bp with a minimum distance of 500 bp from the next TSS or TTS, and the exon has to be surrounded by AG-GT. The default annotation file gencode.v19.annotation.gtf and the reference genome GRCh37.p13.genome.fa were downloaded from https://www.gencodegenes.org/releases/19.html (May 2018) and used for subsequent analyses.

We used three different measurements to quantify splicing ratios (PSI), namely single-cell splicing ratios, pseudo bulk splicing ratios and variance of splicing ratios. To calculate single-cell PSI per cassette exon per cell, we only considered splicing events that were supported by at least five reads and limited the analysis to cassette exons which were observed in at least ten cells. To derive pseudo bulk PSI per cassette exon, we aggregated the single-cell PSI values per cassette exon. The variance of PSI per cassette exon was defined as the standard deviation of PSI across single cells.

### DNA methylation pre-processing and quantification

For DNA methylation data, single-cell bisulfite sequencing (scBS-seq) data was processed as previously described [25]. Reads were trimmed with Trim Galore! [27–29], using default settings for DNA methylation data and additionally removing the first 6 bp. Subsequently, Bismark [34] (v0.16.3) was used to map the bisulfite data to the human reference genome (build 38), in single-end non-directional mode, which was followed by de-duplication and DNA methylation calling using default settings. We removed cells with low alignment rates (alignment rate < 15%) and cells with a library size of less than 1M reads, resulting in 84 iPS cells and 53 endoderm cells with RNA and DNA methylation information.

To mitigate typically low coverage of scBS-seq profiles (20-40%; [17]), we applied DeepCpG [17] to impute unobserved methylation states of individual CpG sites. DNA methylation profiles in iPS and endoderm cells were imputed separately. The cell type specific models were built using CpG and genomic information according to DeepCpG’s set-up of a joint model (see [17] for details and default values; see Table S1 for imputation accuracy as measured on a validation set per sample).

Predicted methylation states were binarized according to DeepCpG probability outputs as follows: sites with a probability of equal to or lower than 0.3 were set to 0 (un-methylated base), all methylation sites with a probability of greater than 0.7 were set to 1 (methylated base). Intermediate methylation levels were handled as missing. After imputation the methylation data was aligned back to human genome version 37 to match the expression data, using the UCSC lift-over tool [35].

We integrated the imputed methylation information into the DNA sequence by distinguishing methylated (‘M’) and un-methylated (‘U’) cytosines. Cytosines without methylation information after imputation were assigned the value of the closest cytosine with methylation information. If there was no methylation information within 900 bp around the cytosine, its state was set to un-methylated.

### Cell and gene model assumptions

To assess if our PSI variation patterns follow the gene or the cell model[18], we compared the distribution of splicing rates to a binomial distribution that is expected according to the cell model, and to the expected distribution according to the gene model.

The cell model assumes that each individual cell expresses only a single splice isoform, and hence models PSI variation as a bimodal distribution at the single cell level. The alternative gene model assumes splicing regulation on the gene level. The mean PSI of a gene is determined by the sequence. Each time a gene is transcribed, the probability of exon inclusion equals mean PSI. However, the limited number of transcripts leads to fluctuation in the observed PSI, and the binomial distribution is restrained by the upper boundary of the standard deviation. To obtain this upper boundary we simulated PSI of each cell as a binomial distribution and calculated the standard deviation across the cells. We only considered genes that were covered by at least five reads per cell in least ten cells. To obtain the mean standard deviation, we repeated this simulation 400 times.

### Sequence features

The genomic features used to predict the splicing ratios and its variance were based on the features described by BRIE and Xiong *et al.* [5, 8]. As these features were specifically designed to study exon skipping events at cassette exons, they are designed to capture sequence variation in the following five genomic contexts: the alternative exon itself, the two neighboring exons and the introns between the exons. Logarithmic length, relative length and the strength of the splice site motifs at the exon-intron boundaries were calculated per genomic context. The strength of the splice site was defined as the similarity between this splice site and known splice motives.

Additional features were calculated on seven genomic contexts, namely the alternative exon itself, the two neighboring exons and the 5’ and 3’ boundaries of the introns. Only the two boundary contexts of the introns (300 bp length) were used since intron length is highly variable and the boundaries are the most relevant contexts for splicing.

Altogether, 607 features were calculated for these seven genomic contexts per cassette exon: PhastCons scores [36] that describe sequence conservation, length of the sequence contexts, and sequence composition based k-mer frequencies (with k ≤ 3) ("genomic" features, Methods, **Table S4**). The k-mers reflect the percentage of nucleotides in the context that match the respective specific motif. The PhastCons scores were retrieved for alignments of 99 vertebrate genomes with the human genome from hg19.100way.phastCons.bw from UCSC (May 2018) [35].

In addition to the genomic features, we defined up to 826 DNA methylation features derived from the imputed DNA methylation information, including an extended k-mer alphabet that takes the methylation status into account, as well as DNA methylation average and variance (across CpG sites), in each of the seven sequence contexts of a cassette exon. Methylation features describe methylation patterns of either individual cells ("genomic & cell methylation" features) or averaged across cells ("genomic & mean methylation" features; **Table S4**). More specifically, for the single-cell PSI model, we considered cell-specific methylation levels; the k-mer features were extended by including un-methylated (‘U’) and methylated (‘M’) cytosine into the alphabet as follows: Cytosines without methylation information after imputation were assigned the value of the closest cytosine with methylation information. If there was no methylation information within 900 bp around the cytosine, its state was set to un-methylated. For the bPSI model, we included the mean frequencies of the k-mers that contained ‘M’ or ‘U’ across cells and the averaged methylation values as described above.

### Splicing categories

In bulk RNA-seq data, splicing events can be broadly categorized into two major categories: included and excluded. Leveraging the single-cell information, we defined more fine-grained splicing categories that reflect both, splicing rates and splicing variability across cells (inspired by Song *et al.*, 2017 [12]): 1) excluded (mean PSI < 0.2), 2) included (mean PSI > 0.8), 3) overdispersed, 4) underdispersed and 5) multimodal (**Fig. 3a**). The later three categories categorize the extent of splicing variation across cells, since cassette exons with intermediate average splicing rates (here 0.2 ≤ mean PSI ≤ 0.8, **Fig. 1**) exhibit substantial differences in splicing variance. To characterize cells into these three categories, we calculated the distribution of the distance between the observed and the expected variation per cell type. The expected variation was calculated by a scaled binomial standard deviation, using: where is the scaling factor and the mean splice rate of the alternative exon [18], fit to all data points. We then defined the overdispersed cassette exons as those for which the deviation from the expected PSI was higher than the 3rd quartile plus 1.5× interquartile range (IQR, corresponding to > 0.016 in iPS and > 0.022 in endoderm). Likewise, for definition of the underdispersed cassette exons, we used the 1st quartile minus 1.5× IQR as threshold (corresponding to < −0.032 in iPS and < −0.039 in endoderm cells). The remaining cassette exons were assigned to the multimodal category.

### Relating DNA methylation heterogeneity and splicing

We applied Spearman correlation to link splicing at a single locus to variation in DNA methylation observed between cells. The test was performed per sequence context of the cassette exon (**Fig. 1c**). We only considered cassette exons where variation in splicing and variation of DNA methylation of the relevant context were observed. In total, 5,280 iPS and 2,622 endoderm cassette exons were tested. The P-values were adjusted for multiple testing using the Q-value [37, 38] package in R. The gene enrichment across the cassette exons was performed using g:Profiler [20] (Version: 2017-10-25, g:Profiler Ensembl 90), using all observed cassette exons per cell type as background. Multiple testing correction for the enrichments was performed within g:Profiler.

### Prediction of PSI and categories

We applied linear ridge regression to model single-cell and pseudo-bulk PSI and (multi-class) logistic ridge regression to model PSI categories. The models are based on only the genomic features or on both, genomic and DNA methylation features. The performance of linear models was evaluated using Pearson r^2^ between predicted and observed splicing rates. For the multi-class prediction models, we applied a one-versus-rest scheme and report the per-category and the macro-average area under the receiver operating curves (AUC). To determine the most relevant individual features, we additionally trained regression models based on each single feature. Per feature we report, in the case of the linear models, Pearson correlation (r, r^2^) and, in the case of the logistic models, the absolute weight multiplied by the standard deviation of the feature, and the AUC. We assessed performance and parameters of models by using a 10-fold cross validation (CV) with fixed training-validation splits. To assess variability of prediction performances, we repeated the CV procedure four times with different CV splits. Error bars indicate plus or minus one standard deviation of the respective statistic (AUC, r^2^).

### Replication cohort

To replicate our results, we processed the mouse ES single-cell scM&T-seq data (n=80) presented in Angermueller *et al.* [14]. We reprocessed the aligned RNA and DNA methylation data to quantify splicing following the same protocols that were applied to the human data. With the following changes: GRCm38 was used as a reference for imputation, genome and transcriptome annotations were based on gencode v18 (“GRCm38.p6.genome.fa” as genomic, “gencode.vM18.annotation.gff3” as transcriptomic reference, available at: https://www.gencodegenes.org/mouse_releases/18.html [August 2018] and conservation scores were taken from “mm10.60way.phastCons.bw” downloaded from UCSC [35] (August 2018).

Out of the 80 cells, in total 12 cells did not pass quality control on the transcriptome data, cells with less than 500,000 reads sequenced and less than 80% of the reads aligned to the genome were removed. Additionally 4 cells did not pass quality on the DNA methylome data, cells with less than 1 million DNA methylation reads aligned and bismark mapping efficiency below 7% where discarded. The filters yielded 68 cells that were used for the main splicing analysis and 64 that are used for the analyses including DNA-methylation data. In these cells we quantified between 649 and 1,433 cassette exons per mouse ES cell (minimum coverage of five reads), in the replication analysis we considered 2,194 exons that were supported by at least ten cells.

### Availability of source code

Python and R were used for data processing, modelling and visualization of the results. Software and scripts are available as jupyter notebooks at https://github.com/PMBio/scmt_splicing. All regression models are based on implementations available in the package scikit-learn [39].

## List of abbreviations

PSI: splicing ratio
iPS cell: induced pluripotent stem cell
ES cell: Embryonic stem cell

## Declarations

### Ethics approval and consent to participate

All samples for the HipSci resource were collected from consented research volunteers recruited from the NIHR Cambridge BioResource (http://www.cambridgebioresource.org.uk). Samples were collected initially under ethics for iPSC derivation (REC Ref: 09/H0304/77, V2 04/01/2013), with later samples collected under a revised consent (REC Ref: 09/H0304/77, V3 15/03/2013).

### Availability of data and material

All sample metadata can be accessed via the HipSci data portal (http://www.hipsci.org), which references to EMBL-EBI archives that are used to store the HipSci data. DNA methylation and RNA expression data will be shared via the European Nucleotide Archive (accession number pending). Replication data is available under bioproject: PRJNA300666 on SRA. The model and data analysis scripts are available as jupyter notebooks at https://github.com/PMBio/scmt_splicing.

### Authors’ contributions

S.M.L.: data processing, splicing ratio modeling, deep modeling, writing of the manuscript L.U.: splicing variation modeling, supervision of splicing modeling and deep modeling, writing of the manuscript

D.J.M.: gene expression data processing, cell-level feature calculations

M.C.: data generation, editing

S.C.: data generation, editing

S.A.: data generation, editing

I.E.: supervision of original master project, editing

L.V.: data generation, editing

W.R.: data generation, editing

O.S.: supervision of modelling, writing of the manuscript, study design

M.J.B.: methylation data processing, supervision of splicing modelling, writing of the manuscript

## Acknowledgements

We thank the staff in the Cellular Genetics and Phenotyping and Sequencing core facilities at the Wellcome Trust Sanger Institute. We acknowledge the participation of all NIHR Cambridge BioResource volunteers, and thank the NIHR Cambridge BioResource centre staff for their contribution. We thank the National Institute for Health Research and NHS Blood and Transplant. The NIHR/Wellcome Trust Cambridge Clinical Research Facility supported the volunteer recruitment. We thank Y. Huang for helpful discussions and comments on the manuscript.

